# Hippocampal State Transitions at the Boundaries between Trial Epochs

**DOI:** 10.1101/443077

**Authors:** David A. Bulkin, David G. Sinclair, L. Matthew Law, David M. Smith

## Abstract

The hippocampus encodes distinct environmental and behavioral contexts with unique patterns of activity. Representational shifts with changes in the context, referred to as remapping, have been extensively studied. However, less is known about the nature of transitions between representations. In this study, we leverage a large dataset of 2056 neurons recorded while rats performed an olfactory memory task with a predictable temporal structure involving trials and inter-trial intervals, separated by salient boundaries at the trial start and trial end. We found that trial epochs were associated with stable hippocampal population representations, despite moment to moment variability in stimuli and behavior. Representations of trial and inter-trial interval epochs were far more distinct than spatial factors would predict and the transitions between the two were abrupt, with a sharp boundary suggestive of a dynamic shift in the representational state. This boundary was associated with a large spike in multi-unit activity, with many individual cells specifically active at the start or end of each trial. Both epochs and boundaries were encoded by hippocampal populations, and these representations carried information on orthogonal axes readily identified using principal component analysis. We suggest that the activity spike at trial boundaries might serve to drive hippocampal activity from one stable state to another, and may play a role in segmenting continuous experience into discrete episodic memories.

## Introduction

For animals to successfully interact with their environment they need to construct neural representations that allow them to identify the current context and select appropriate behavioral responses, and they need to rapidly transition between these representations when the context changes. A large literature has suggested that hippocampal activity patterns represent the environmental context (Colgin et al., 2008; Holland and Bouton, 1999; Nadel, 2008; Rudy, 2009) and we have shown that distinct hippocampal representations are essential for the ability to retrieve context-appropriate memories while avoiding interference from memories that belong to other contexts (Bulkin et al., 2016; Butterly et al., 2012; Smith and Bulkin, 2014). The hippocampus has the capacity to generate many distinct representations (Alme et al., 2014) and the environmental factors that induce the formation of a new representation (i.e. remapping) have been studied extensively (Anderson and Jeffery, 2003; Colgin et al., 2008; Leutgeb et al., 2007, 2005; Muller and Kubie, 1987; Schlesiger et al., 2018). Changes in the non-spatial characteristics of the context, such as behavioral demands, strategy and motivation, are also known to induce remapping (Eschenko and Mizumori, 2007; Ferbinteanu and Shapiro, 2003; P. J. Kennedy and Shapiro, 2009; Skaggs and McNaughton, 1998; Smith and Mizumori, 2006; Terrazas et al., 2005; Wood et al., 2000). Studies of hippocampal representations during context transitions have shown patterns that are rapid and abrupt, rather than a gradual progression through intermediate representations (Jezek et al., 2011; Kelemen and Fenton, 2010; T. J. Wills et al., 2005). However, these studies necessarily involved artificial experimental conditions unlike those commonly encountered in day to day experience (e.g. an unexpected and dramatic change in the visual environment, described as ‘teleportation’ by the authors (Jezek et al., 2011)). Much less is known about more mundane and highly predictable changes in the context such as walking from the living room into the kitchen. Indeed, it is not clear whether the hippocampus treats contiguous spaces as distinct contexts and if so, how hippocampal representations transition from one to the other.

In this paper we leverage a large dataset of 2056 neurons recorded during a complex multi-stimulus olfactory discrimination task with two behaviorally and spatially distinct areas (a trial area and an inter-trial waiting area) and a predictable trial structure to interrogate the dynamics of hippocampal representations. We discovered that hippocampal populations form two distinct representations of the trial and ITI epochs, and that the shift between these representations was accompanied by a surge of activity among subsets of hippocampal neurons. These firing patterns resembled a phase transition: the hippocampal state before trials transformed to a distinct state during trials, and then transformed back at the end of trials, with an identifiable transitory activity pattern between states.

## Results

We recorded the responses of hippocampal CA1 neurons while rats were engaged in a memory-guided odor discrimination task (Butterly et al., 2012). On each trial, a removable divider was lifted and rats ran from an inter-trial waiting area to approach two cups containing scented digging medium (Fig.1A). Odors were drawn from a set of 16 distinct odors, presented in 8 pairings with one odor in each pair always rewarded with a buried sucrose pellet. On some trials (Fig. 1B, E), rats approached the rewarded cup and dug for a reward. On other trials rats approached the unbaited cup first (Fig. 1F), and either correctly rejected the stimulus (Fig. 1C) or incorrectly dug for a reward (Fig. 1D) after which the trial continued until they obtained the reward in the baited cup. Following the reward, rats returned to the inter-trial waiting area. Recordings took place over a period of up to 10 days, as rats learned reward contingencies for two sequentially presented sets of odor pairings in a task design used to probe for mnemonic interference. Results of the investigation into interference have been reported previously (Bulkin et al., 2016).

**Figure 1.**
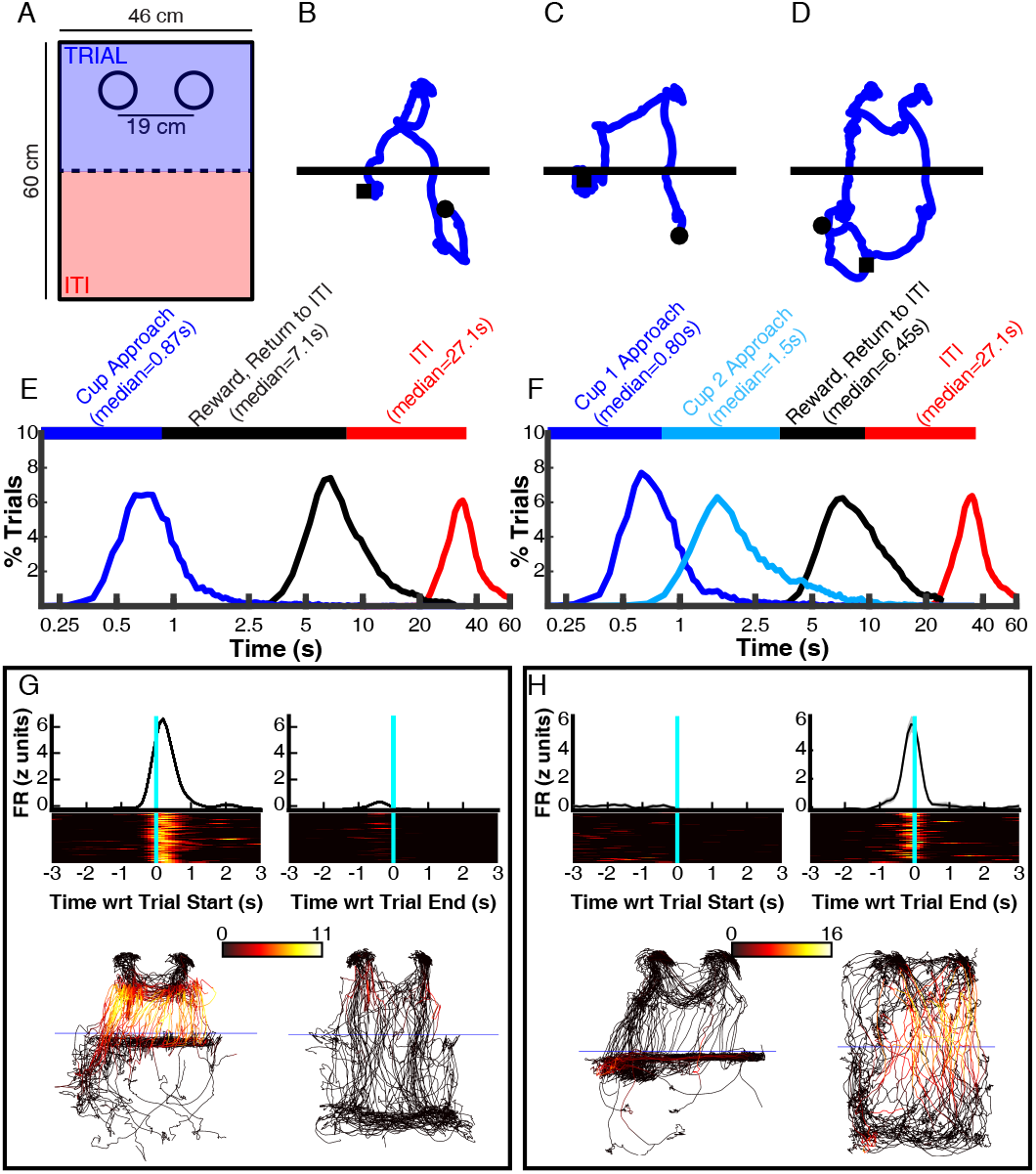
Reliable neuronal responses to the start and end of trials despite large variability in spatiotemporal patterns of behavior. Rats performed trials in 60cm × 46cm wooden boxes (A). The boxes were bisected by a removable divider (dotted line in A). One side of the box served as the inter-trial waiting area, and rats performed trials in the other side. During the trials, rats dug for a reward buried in one of two cups, placed in cup holders indicated by the two circles in A. Panels B through D show example positional trajectories, beginning 3 seconds before the trial start (black squares) and extending 3 seconds after the trial end (black circles). B shows a trial where the rat first approached the cup on the right, which was the baited cup for this trial, and dug for a reward. C shows a trial where the rat approached and sampled the unbaited cup (on the left), correctly rejected it, and then dug for a reward in the baited cup on the right. D is similar to C, except that the rat made an error by digging in the cup on the left, but then approached the baited cup on the right and found the reward. Panels E and F illustrate the time line and variability in how the trials proceeded. The median time of arrival at the cup (blue), the time spent retrieving the reward and returning to the ITI side of the box (black), and the duration of the ITI (red) are given. The histograms illustrate the variability in the duration of these epochs. Panel F shows the same data for trials in which the rats approached both cups (i.e. trials with trajectories like those shown in C and D). Note the use of a log scale for the abscissa in E and F. Panels G and H show data from an example trial start and trial end neuron. At the top the average firing aligned on the start (left) and end (right) is shown. Below a heatmap shows the firing on each trial. At the bottom, the trajectory of the rat is shown for a period matching the above plots, with color indicating the instantaneous firing rate of the neurons.

### Individual Neurons Respond at Trial Boundaries

Many neurons showed transient increases in activity at the start and end of trials, often selective for one of these two epochs. Figure 1G-H show the responses of example ‘trial start’ and ‘trial end’ neurons. The upper panels show responses that are strongly time-locked to the start and end of trials, despite considerable variability in timing of behavior (Fig. 1E-F). The lower panels of Figure 1G-H show the trajectory of the rat with color indicating the instantaneous firing rate. These data show that elevation in activity was spatially unrestricted, and that the rat often traversed similar territory during these two epochs, suggesting that activity was better explained by a temporal description of firing rate than by a spatial account.

The large numbers of neurons with activity patterns like those seen in Figure 1G-H led to transient increases in multi-unit firing rate that began just before the start of trials (as the divider was lifted) and then again at the end of the trials as rats returned to the inter-trial waiting area (Fig. 2A-D). Similar to the example responses shown in Figure 1G-H, this activity was not attributable to purely spatial factors. Most neurons were selective for either the trial start or trial end (Fig. 2E, blue and red) even though these epochs occurred in similar locations. Firing was not distributed randomly around the environment, but instead we observed elevated activity near the boundary between the trial and ITI areas (Fig. 2F). These apparent ‘spatial’ regions of elevated firing were clearly modulated by the start and end of trials. Firing measured at the same locations during epochs associated with the trial start and end was distinct (Fig. 2G, red and blue traces), and was elevated compared to firing far from trial boundaries (Fig. 2G, black trace), indicating that activity was not modulated solely by spatial location.

**Figure 2.**
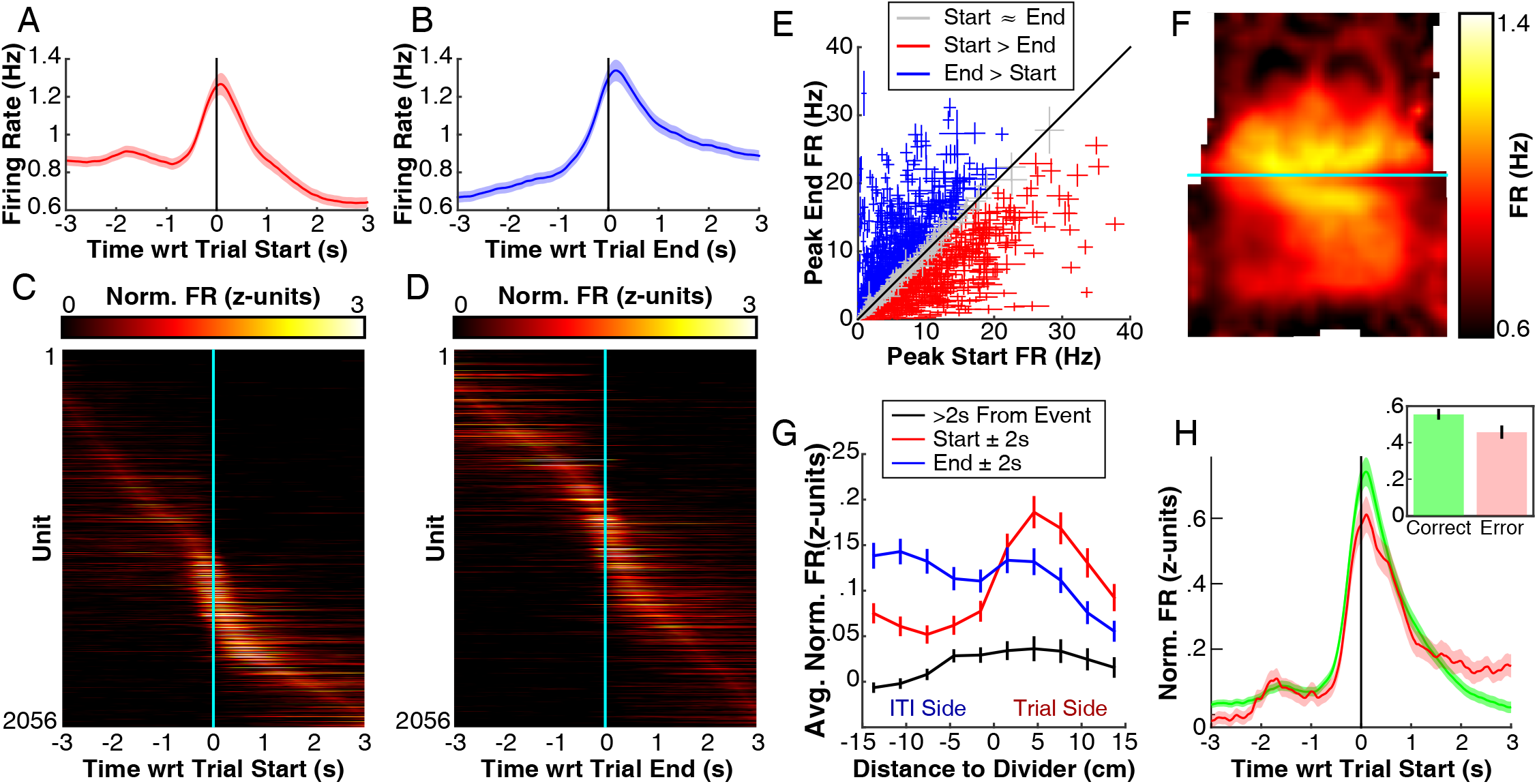
Increase in activity of hippocampal neurons at trial start and end. Panels A and B show average multi-unit firing rate aligned on trial start and end. Firing rates of each unit were binned (100ms bins) and smoothed with a 5 bin moving average. The shaded region indicates SEM over units. Panels C and D show average normalized firing rate aligned on trial start and end for all units. Binned activity was normalized by z-scoring using the average and standard deviation of each unit’s rate over the entire session. The trial-averaged traces were then sorted based on the time of the maximum rate. Panel E shows a scatter plot of peak firing rate of each neuron in a window +/- 3s with respect to trial start (abscissa) versus trial end (ordinate). For each unit a single crosshair is plotted, centered on the average peak rate, and extending +/- one SEM (over trials). Units with SEM overlapping with unity are shown in gray (513/2056 units), those above unity are shown in blue (810/2056 units), and those that fall below unity are shown in red (733/2056 units). Panel F shows the average firing rate in 1.5 cm square spatial bins. Points between the centers of the bins have been linearly interpolated. The horizontal line indicates the location of the removable divider (see Fig. 1A). Panel G shows average multi-unit firing rate for locations in 3cm bins in the axis orthogonal to the divider (i.e. vertical in F), calculated separately for rates occurring within 2 seconds of the start of the trial (red) or end of the trial (blue) or more than 2 seconds from either (black). Panel H shows average firing rate around the start of the trial plotted separately for correct (green) and error (pink) trials, only neurons with trial start responses have been included (688 neurons, Fig. 3A). The inset shows the average activity in a window +/-500ms on the trial start. Shaded region and error bars indicate SEM over units.

Transient activity at the start and end of the trials was not likely associated with sharp-wave ripples. Rats were rarely immobile at these times, and field potentials showed robust theta oscillations, and so ripples were infrequent at the start and end of trials (Buzsáki et al., 1992). Increased activity at the trial start/end was also not attributable to increased running speed at the start and end of trials. Although activity was correlated with running speed overall, we found that firing rates were higher at the time of the trial start and trial end than at instances of similar running speed occurring during the trial and ITI epochs, suggesting that trial start/end activity was above what would be expected based on running speed alone (Supplemental Fig. 1). To statistically confirm this, we computed a series of linear regressions for each session defining multi-unit firing rate as a function of running speed, separately for data selected from the trial start and trial end and the inter-trial interval (ITI). The slopes of the resulting regression lines taken from the trial start/end data were similar to the slopes based on ITI data (paired t-test, p>.01 for start and end), yet the intercept was significantly higher for data selected from the trial start (paired t-test, p<10^−6^) and end (paired t-test, p<10^−6^). This suggests that activity at these times showed a global increase unrelated to running speed.

Interestingly, the magnitude of the trial start response (average firing rate +/- 500ms around trial start) was somewhat larger on trials in which the rat made the correct choice on the subsequent trial (Fig. 2H; 641units, paired t-test: p<.001). This effect was probably not driven by reward related activity, or a reduction in the reward prediction error when rats identified the odor and could therefor predict an impending reward. Rats rarely arrived at the first cup within 500ms (Fig. 1E-F), and we saw no evidence of a decision (e.g. a change in trajectory) before this time.

To assess the relative contributions of spatial and temporal factors in shaping the activity of individual neurons, we modeled the firing rate of each neuron as a function of the rat’s location and the time of the nearest trial start and end. Because position and time partially covaried (i.e. position was not random at times near the trial start and end), we used an extension of a constrained Poisson generalized linear model (GLM) that orthogonalizes covariates to disambiguate the independent contributions of factors affecting firing rate (Truccolo et al., 2004). This strategy has been previously used specifically for distinguishing effects in the face of nuisance correlations between factors shaping hippocampal activity (Lepage et al., 2012; MacDonald et al., 2011). For each neuron we fit three Gaussian functions to the average activity: two one dimensional curves that described firing rate as a function of time with respect to trial start and end, and a two dimensional surface that described firing rate as a function of the location of the rat. We then described the firing rate of the neuron as the weighted sum of these three functions (effects of trial start, trial end, and position), using the projection described in Lepage et al (2012) to form estimates of temporal effects that could not be accounted for by covariation between the times of trials and the location of the rat. Finally, we computed the statistical significance of each coefficient via a normal approximation of the bootstrap estimate (Efron and Tibshirani, 1994). Supplementary Figure 2A-F show responses and fits of example temporally and spatially modulated neurons that were disambiguated by this analysis.

The majority of neurons were significantly modulated by space (1601/2056; Fig. 3A). Yet many of these neurons showed additional modulation by the trial start or end (859/1601; Bonferroni corrected for tests of trial start and end responses). Importantly, because the model orthogonalized spatial and temporal covariates, the trial start/end responses were not spuriously identified due to rats traversing through a place field at the beginning or end of trials. Rather, neurons represented both location and time with respect to the trial boundaries. Approximately half of the neurons showed some modulation by one of the temporal factors (1004/2056, Bonferroni corrected), and a similar quantity of cells had activity that was affected by the trial start and trial end (start: 688 neurons; end: 757 neurons). Plotting the average firing rate for start/end responsive and spatially sensitive neurons revealed that firing patterns were similar for neurons with a trial start/end response whether or not the neuron was also modulated by space, and that spatially sensitive neurons that lacked a trial start/end response showed larger firing rates during ITI epochs (Fig. 3B-C; Supplemental Fig. 2G-H). In order to control for the possible contribution of running speed to trial start/end firing, we repeated the GLM and included a linear term for running speed. Firing was modulated by running speed for many neurons (773/2056), but accounting for variance due to running speed had little effect on the number of neurons marked as responsive to the trial start/end (Supplemental Fig. 2I-J).

**Figure 3.**
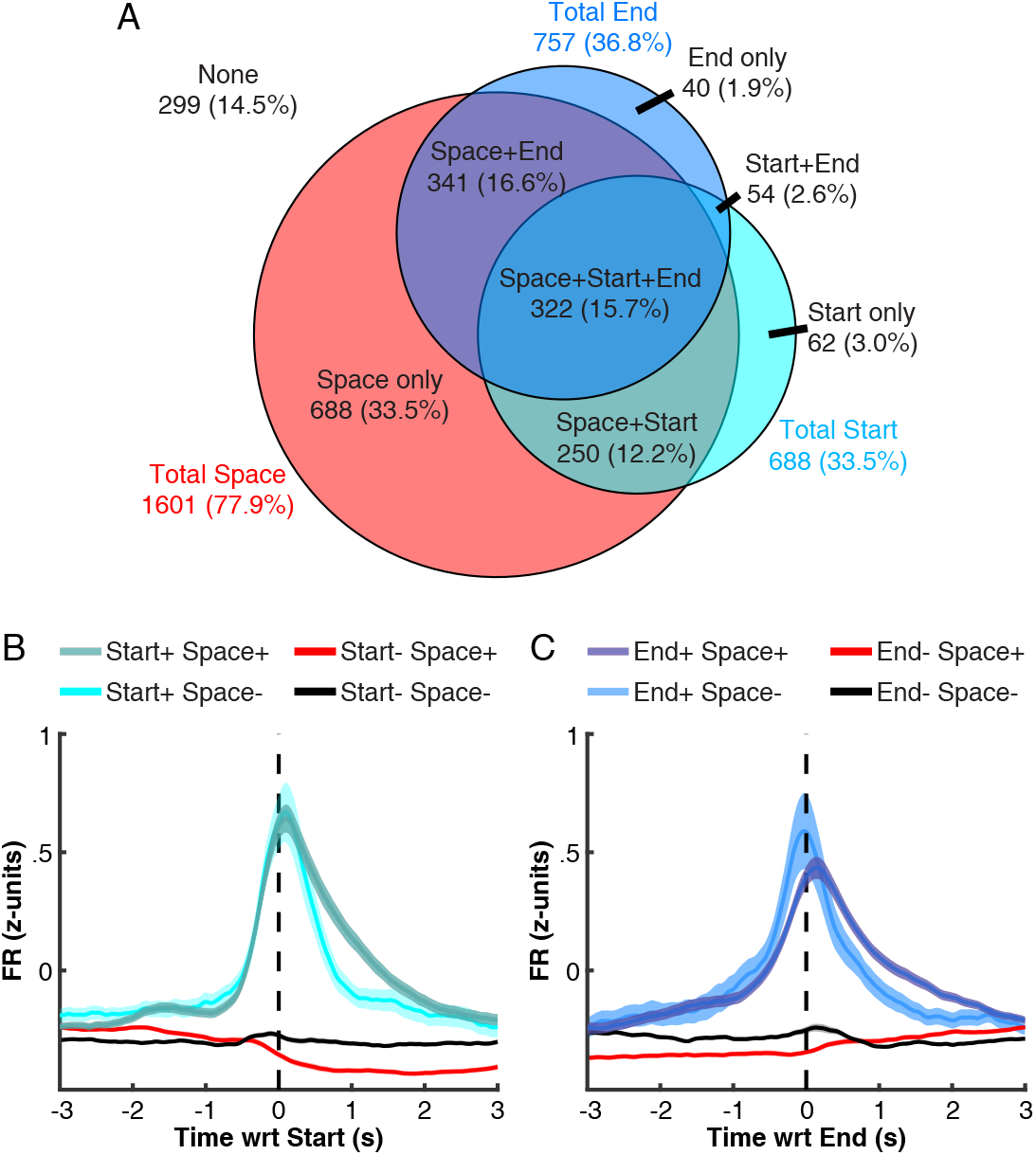
Independent Spatial and Temporal Responses in Overlapping Neuronal Populations. Panel A shows a Venn diagram indicating the classified responses of neurons. Each unit was submitted to a generalized linear model (GLM) which orthogonalized components defined by spatial and temporal Gaussian fits to average response data. The diagram tallies the number/percent of neurons with a significant term in the model for the noted component. Panels B and C show average normalized firing rate for units with (+) and without (-) significant spatial and trial start (B) or trial end (C) coefficients. The GLM successfully identified start and end responsive neurons, evidenced by the clear peak in the average firing rate of these neurons compared with unresponsive neurons. Units classified as exclusively spatial (i.e. Start-/End- Space+) showed somewhat elevated firing rates in the inter-trial interval (before the trial start in B and after the trial end in C).

### Distinct Population States Represent Trial and ITI Epochs

Inspection of the neural activity in Figure 2 revealed two important details about the dynamics of hippocampal firing as rats started and ended trials: a prominent increase in firing at the trial boundaries (Fig. 2A-B), and distinct populations of neurons were recruited during trial and ITI epochs (Fig. 2 C-D). Unlike the transient activity increases at the trial boundaries, non-overlapping populations during trials and ITIs are potentially consistent with spatial models of hippocampal activity as these two epochs were necessarily in distinct spatial locations (Fig. 1A). This presents an essential challenge of comparing temporal and spatial dynamics in the hippocampus: distinct events often occur in distinct locations. We next performed a series of population analyses to understand whether the extent of neural dissimilarity between trials and ITIs could be explained based on differences in the rat’s position, or whether trial boundaries marked a state transition such that neuronal dissimilarity exceeded what would be expected based on positional disparity alone.

To measure ensemble similarity, we tabulated population vectors (PVs; vectors containing firing rates in 100ms bins) and computed pairwise correlations between them. We averaged the pairwise correlation coefficients between PVs drawn from different trials, at times surrounding the start and end of trials (+/- 3s). Figures 4A-B show the average values (across recording sessions) for each pair of comparisons. The color of each point in the images indicates the average cross-trial correlation between one temporal bin and another. Points lying on unity quantify the similarity of ensemble activity from trial to trial at the same time with respect to the trial start/end, while off-unity points indicate cross-trial similarity at proximal times.

Examination of the cross trial correlation plots (Fig. 4A-B) reveals several striking characteristics of large scale hippocampal activity patterns. Both plots show a strong peak at the center, indicating that trial start/end firing patterns are similar from one trial to the next, an outcome which reflects the reliable bursts of firing seen at the trial boundaries (Fig. 2A-B). Correlations along the unity line declined from this peak as time passed from the trial start, but remained high throughout the trial epoch. For example, firing patterns occurring ~1.5 sec into the trials (see ●, Fig. 4A) were surprisingly well correlated across trials. At that time, the rats had typically arrived at the first odor cup (Fig. 1E-F), encountered one of the sixteen possible odor cues, and decided whether to dig or proceed to the second cup, depending on the valence of the odor cue. Even more noteworthy, firing patterns taken from quite distant times within the trials were also well correlated. For example, the firing patterns occurring 0.5 sec into the trial were surprisingly similar to firing occurring two seconds later (i.e. 2.5 sec into the trial, see ♦, Fig. 4A), despite the fact that the rats were engaged markedly different and highly variable behaviors at those two time points (Fig. 1E-F). Rats were nearly always approaching the first odor cup at 0.5 sec, but at the 2.5 sec time point they could be digging in the first cup, investigating or digging in the second cup (if the first cup was not rewarded), consuming the reward, or returning to the ITI side of the chamber. This was not likely driven by spatial location since the trial structure meant that rats rarely occupied the same location at these two different time points (Fig. 1B-D). Firing patterns within the ITI epoch also showed a large degree of similarity (Fig. 4A lower left and 4B upper right), although these correlations were significantly lower than those for population vectors taken from the trial epoch (Fig. 4C; paired t-test: T(83)=4.59, p<10-4). Another important feature that is apparent in these plots is the sharp boundary between the trial and ITI epochs. In contrast to the remarkable self-similarity of firing patterns taken from within an epoch, correlations were significantly lower for vectors drawn from different epochs (Fig. 4C; paired t-tests: ITI vs. cross T(83)=18.89, p<10-31; trial vs. cross T(83)=14.27, p<10-23). This suggests that activity was distinct across the two epochs (Fig. 4A-B, upper left and lower right). Indeed, correlations for vectors taken only 1 sec apart but from different epochs (i.e. 0.5 sec before and 0.5 sec after trial start, see □, Fig.4A) were much lower than those for vectors taken twice as far apart but within the trial epoch (see ♦, Fig. 4A). Similar patterns were found when correlations were measured using Kendall’s rank correlation coefficient, which is arguably more robust to the relatively sparse firing patterns seen in hippocampus (Neymotin et al., 2017) (Supplementary Fig. 3A-C).

**Figure 4.**
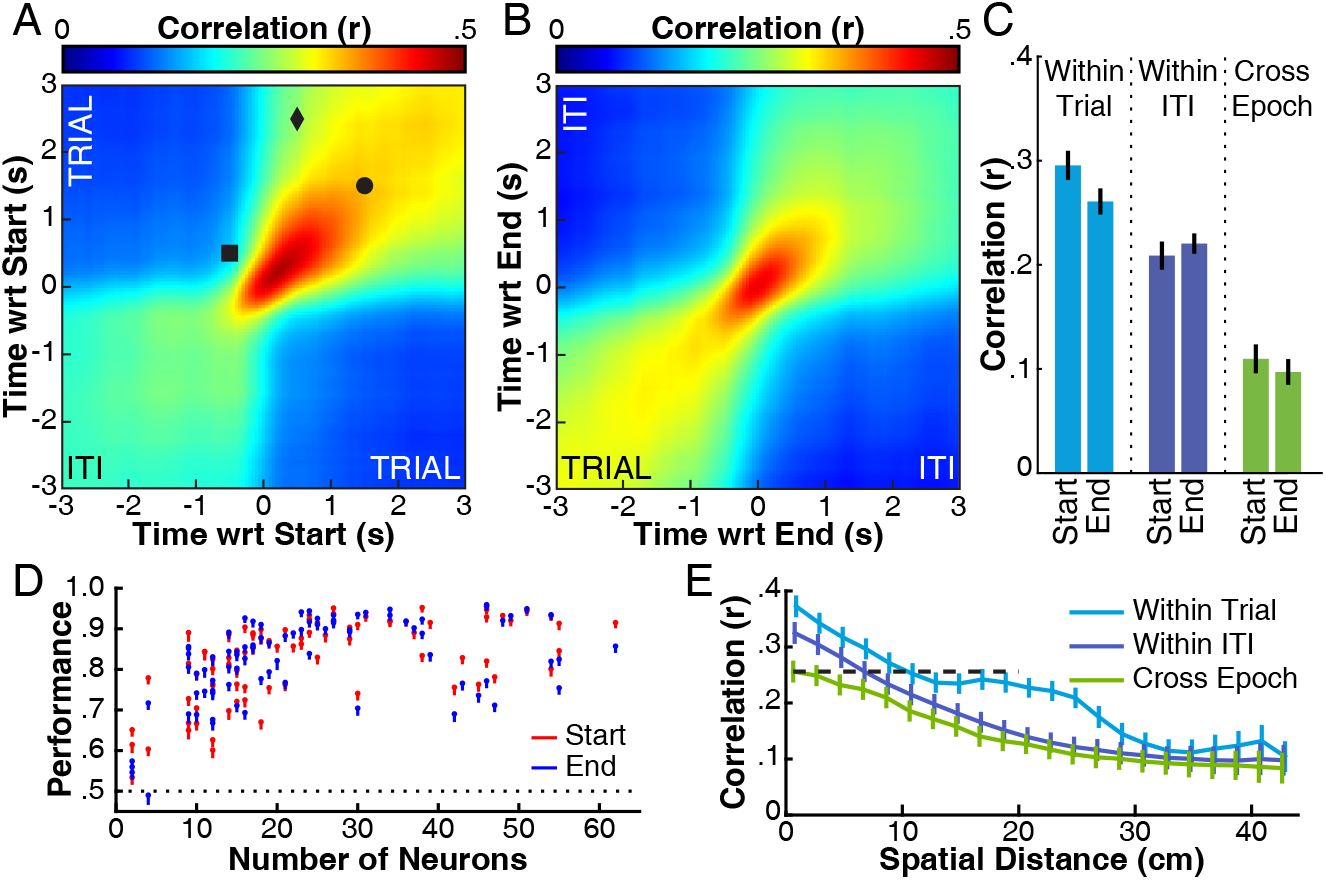
Distinct population states in trials and inter-trial intervals. Panels A and B show cross trial population vector correlations around trial start and end. For each session, firing rates were tabulated to form population vectors and pairwise correlations were computed using vectors in a 6 second window around the trial start (A) or end (B) from different trials (each pairwise correlation involved two unique trials at two time points). Correlation values were averaged to form a map for each recording session, the average of these maps is shown. Points in the image between bins have been linearly interpolated. The symbols overlaid on the plot in A highlight times of interest discussed in the text. Panel C shows a summary of cross-trial correlations. The height of the bars indicates the average value in the corresponding quadrant of the maps shown in A-B. Error bars indicate SEM across sessions. Panel D measures the performance of a linear discriminant classifier trained with a subset (50%) of population vectors selected from trials and ITIs to classify the epoch of the vector (i.e. whether it occurred in an ITI or a trial). The classifier was tested separately on vectors selected from a window 3 seconds before/after the trial start (red points) and trial end (blue points). Confidence intervals were estimated using an iterative process, randomly selecting vectors 1000 times, with the 5% lower C.I. identified as the 5^th^ percentile of the iterated dataset. Lines span from this point to the median performance across iterations for each session. The abscissa indicates the number of simultaneously recorded neurons in the session. Panel E shows pairwise correlations between population vectors as a function of the distance between positions that the rat occupied when the activity occurred. Correlations were computed separately for pairs in which both vectors were selected from trial epochs (cyan), ITI epochs (blue), or when one vector was selected from each epoch (green). Error bars indicate SEM.

The striking self-similarity of firing patterns within each epoch, and the sharp decline in similarity when rats transitioned from the trial epoch to the ITI, suggest that hippocampus treats these two epochs as distinct contexts. Consistent with this idea, we could accurately decode population activity as belonging to the trial or ITI using linear discriminant analysis. We used an iterative process wherein we trained linear classifiers using a randomly selected subset of half of the PVs and measured performance of the classifiers on the remaining half. We repeated this process 1000 times, providing a distribution of performance values (proportion of PVs correctly classified). Classifier performance (Fig. 4D) was virtually always above chance, and showed high accuracy: on average 85% of vectors surrounding the trial start and 84% of vectors surrounding the trial end were correctly classified. Classification errors were most likely to occur near the trial boundaries (Supplementary Fig. 3F-G), a period when ensembles encoded the boundary itself rather than the surrounding epoch.

While hippocampal output reliably differentiated the task epochs, the increased similarity of ensemble activity within epochs could simply be due to spatial factors since any two PVs drawn from a single epoch were more likely to correspond nearby locations than two PVs selected from different epochs. Even the greater similarity within trials than within the ITI could have been influenced by spatial factors since spatial behavior was more constrained during trials. In order to determine whether these spatial factors did, in fact, account for the within-epoch similarity, we compared PV correlations for subsets of data with fixed ranges of spatial distance. We labeled each pair of PVs using the distance between the associated positions (locations occupied by the rat at the time of the PV) and binned pairwise correlation values based on distances (Fig. 3E). If ensemble activity was governed purely by space, the selected epoch would make no difference in the correlation values and an overall decrease in correlation with distance would be expected. In fact, even at very low distances, PVs were more similar within epochs than across epochs, and within trial similarity was higher than within ITI similarity (repeated measures ANOVA: main effect of epoch F(2,166)=19.15, p<10-7; main effect of distance F(21,1743)=182.9, p<10-15; interaction F(42,3486)=6.053, p<10-15). Indeed, correlations for population vectors taken 10-20 cm apart during a trial were as similar as population vectors taken just 1 cm apart but which spanned the trial start boundary (Fig. 4E, dashed line). This increased similarity between vectors selected from trial epochs persisted at larger distances, with a noticeable ‘bump’ in the similarity curve for PVs that occurred when positions were separated by about 20cm. This distance is of particular note as the odor stimuli used in the experiment were presented in cups separated by 20cm (Fig. 1A) and we occasionally observed examples of individual neurons that showed increased activity as rats sampled the odors and dug in the cups, regardless of which cup (see also (Eichenbaum et al., 1987; Muzzio et al., 2009)). The observation that the trial and ITI epochs were more similar than would be suggested by spatial considerations alone is consistent with the idea that the hippocampus represents the two epochs as distinct contexts and differentiates them accordingly.

### Dynamics of Hippocampal Ensembles during State Transitions

The dissimilarity between population activity in trial and ITI epochs, above what is predicted from space alone, suggests that hippocampal ensembles undergo a comprehensive state transition at the boundaries of each trial event. Under a phase transition framework, the increase in hippocampal multiunit activity at the start and end of trials might serve to drive this transition, pushing the hippocampal state past a critical point to allow a shift in representational state (Steyn-Ross et al., 2010). As such, we next sought to characterize the hippocampal state itself rather than relying on pairwise correlations to make inferences about the clustering of hippocampal representations. We took a dimension reduction strategy using principal components analysis (PCA). Because PCA produces an orthogonal transformation to a set of linearly uncorrelated variables accounting for descending quantities of variance, it allows for a representation of high dimensional neural activity that captures important covariation among ensembles. However, while the individual components identify a mapping of the raw data based on variance, the sign of PC scores is irrelevant. Thus, averaging PC scores across sessions provides no information on how a typical ensemble changes. To circumvent this issue we built a large matrix of pseudo-population vectors containing the firing rates of all of the units in our dataset (n=2056). To combine PVs across multiple sessions, we labeled vectors based on their time relative to the trial start and trial end. We took activity from 100ms bins extending +/-5 seconds around each trial’s start and end. We then randomly sampled (with replacement) a vector from a given bin from each session 200 times, generating typical 2056 dimensional vectors for that moment in time. Repeating this process across bins produced a matrix with 40000 observations (200 time points x 200 iterations). We subjected this entire matrix to PCA. Importantly, although the pseudo population vector matrix was assembled using temporal labels, PCA is blind to these labels and simply provides loadings (i.e. a coefficient for each neuron) such that the first principal component accounts for maximal variance and each additional component accounts for a decreasing amount of variance.

Figure 5A shows the scores of the first three principal components for each vector in the pseudo-population vector. Points taken from the ITI (before trial start or after trial end) are shown in cooler colors, and points during trials (after trial start or before trial end) are shown in warmer colors. A curve showing the trajectory through PC-space was constructed by applying the coefficients identified from PCA back to the (raw) average firing rates in the time +/- 5s around the trial start and end. The projections of this three-dimensional representation to each of the two-dimensional planes are shown as ‘shadows’ on the axes. PC1, capturing the largest amount of variance, distinguished the epochs: trials and ITIs formed completely non-overlapping clusters (Fig. 5B; see blue vs. red clusters in 5A). PC2 identified trial boundaries, clearly distinct from the trial and ITI epochs, but not from each other (Fig. 5C). PC3 made this distinction, differentiating trial start activity from trial end activity (Fig. 5D).

**Figure 5.**
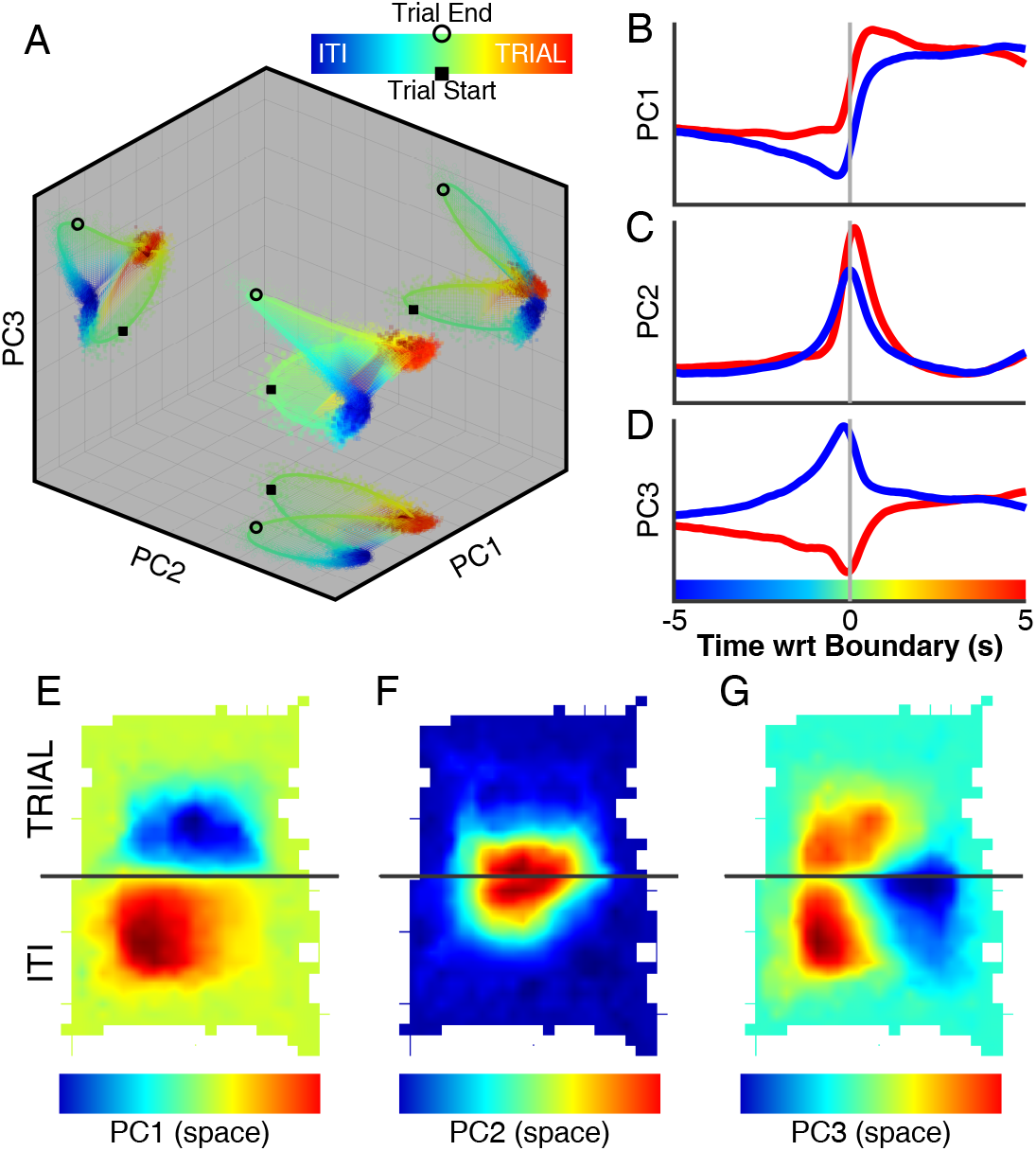
Principal component analysis indicates a phase transition at event boundaries. Using a bootstrap approach, a pseudo-population vector matrix was assembled to simulate typical firing vectors at time points +/- 5 seconds around the trial start and end. This matrix was submitted to PCA to obtain principal component scores for each time sample. (A) Three dimensional plot of the first three principal component scores. Colored squares and circles indicate scores of the principal components around the trial start and end respectively, warmer colors indicate points selected from time bins in the trial while cooler colors indicate time points selected from the ITI. The event boundaries are indicated with black markers. A line traces the trajectory through PC space, computed by applying the coefficients obtained by PCA and taking the weighted mean of average peri-event firing (Fig. 1C-D). The colored surface is shown to aid interpretation of the three-dimensional structure, and was formed by linear interpolation. Projections of the three-dimensional data into each two-dimensional pair are shown on the axis boundaries. (B-D) The PC data shown in A, plotted as a function of time for each principal component. As with the line shown in A, these values were computed by applying the coefficients obtained from PCA to the average firing rate traces for each unit. (E-G) Spatial PC heatmaps computed using the same strategy as in panel A, but assembling a pseudo-population matrix based on spatial location rather than the time of individual population vectors (see text). Coefficients from PCA were applied to the individual unit spatial firing heatmaps to compute a weighted average. The maps have been linearly interpolated between sampled locations.

These results provide a view of the state transition of hippocampal ensembles over the course of trials. Despite a variety of individual neural firing patterns in the trial and ITI epochs, clear clusters of ensemble activity form that identify these distinct contexts. At the start and end of trials, ensembles must transition from one representation to the other, and they do this by traversing orthogonally through the trial start and trial end PC space. This pattern is similar to a phase transition in that one steady state of population activity moves to another state indirectly: the ensemble passes through a specific and reliable intermediary.

To confirm that this approach yielded a view of the hippocampal state that was not artificially imposed by grouping vectors based on time with respect to events, we repeated the analysis but combined vectors across sessions using a purely spatial method. To do this, we formed a matrix of PVs for each session, and labeled each vector with the position of the rat at the time associated with the activity. We then sampled PVs from each session (with replacement), concatenating vectors that occurred when the rat was in the same spatial bin (3 cm square bins). In this manner, we formed a large matrix of vectors in 2056 dimensional space, each vector marking typical population firing rates for a particular spatial location. This is an identical procedure to the method described above but here vectors were combined based on spatial location of the rat rather than the time with respect to trial start and end. We subjected the spatial pseudo-population vector to PCA to obtain coefficients for each neuron, and used the coefficients to create average spatial maps in PC space (Fig. 5E-G). The pattern is strikingly similar to what we found with our time locked analysis. PC1 distinguished the trial and ITI epochs, it was most distinct between regions associated with trials and ITIs (compare Fig. 5E top and bottom). PC2 marked the event boundaries, values near the divider are distinct from values far from the divider (Fig. 5F). In contrast to the time locked analysis where PC3 distinguished the trial start and end (Fig. 5D), the spatially binned PCA did not clearly distinguish them (Fig. 5G). This was expected because the trial start and end occurred in the overlapping locations.

The strong hippocampal transitions at trial boundaries were specifically dependent on trial start/end responsive neurons. When the same analysis was restricted to the subset of neurons with significant event boundary responses, as identified with our GLM approach (Fig. 3), the shape of the resulting pattern in PC-space was virtually identical (Supplementary Fig. 4A). Yet the pattern was completely different when the neurons without trial start/end responses were analyzed (Supplementary Fig. 4B), despite the similarity in sample size (1004 responsive vs. 1052 un-responsive cells). The separation between ITI and trial PVs captured by PC1 was preserved when trial start/end cells were excluded, but the transitional signal between these events that was evident in PC2 and PC3 was completely eliminated.

## Discussion

In this paper we examined the responses of 2056 neurons while rats engaged in an olfactory memory task with a repeated trial structure. We focused our analyses on activity differences between trials and intertrial intervals, as well as the transition between these epochs. We observed a large-scale shift in the activity state of the hippocampus at the start and end of trials, a transient increase in activity that marked a transition between two highly distinct states. We found that individual neurons were modulated by both spatial and temporal factors, driving an ensemble representation that clearly identified the trial and inter-trial interval epochs and the boundaries between them (trial start and trial end). The dynamics resembled a phase-transition-like pattern: populations transformed from one steady state to another at the moment of trial start or end, with a transient increase in firing that co-occurred with the transition.

The self-similarity of hippocampal firing patterns from one trial to another was striking. Although different trials shared some behavioral and sensory features (an increase in running speed, investigation of odors, digging for and consumption of the reward), there was also a great deal of variation from one trial to the next. This included the trajectory of the rat, the olfactory experience of sixteen distinct odors, the left or right position of the reward, and whether the rewarded odor was encountered first by chance or the initial odor cup was rejected in favor of the second. Thus, the similarity of firing patterns was not simply driven by identical sensory input and motor behavior on each trial. Because the sequence of events in the trials was determined by the voluntary behavior of the rat and the randomization procedures (e.g. the left or right location of the rewarded odor cup), the rat’s experience became increasingly distinct as the trial progressed. Despite this increase in behavioral/environmental variability, firing patterns remained quite similar. This is even more striking for correlations of time points that were several seconds apart, when behavior and sensory experience were virtually always different (Fig. 4A). These results are consistent with previous findings that hippocampal firing patterns occupy a local minimum in state space where firing patterns are relatively stable and insensitive to small changes in the environment, until environmental change is sufficient to abruptly push the firing patterns into a new state space (T. J. Wills et al., 2005). However, previous studies involved subtle changes in the shape of the environment and similar foraging behaviors. Here we show that a self-similar hippocampal state persists even in the face of highly variable sensory input and behaviors during the performance of a complex memory task.

The observation of an abrupt shift in the representation when the rats transitioned between the trials and ITIs is similar to findings from studies that manipulated the environmental context (Jezek et al., 2011; Kelemen and Fenton, 2010; T. J. Wills et al., 2005), suggesting that the hippocampus treated the trial and ITI areas as distinct contexts even though they were both part of a contiguous environment which was only divided by a barrier for part of the time. Consistent with this idea, firing patterns for different locations were far more distinct than would be expected based on spatial distance alone as long as they came from different trial and ITI epochs, even for immediately adjacent locations (Fig. 3E). This may be due to the markedly different task demands and motivational characteristics of the trial and ITI epochs since changes in the behavioral context are also known to induce remapping (Eschenko and Mizumori, 2007; Griffin et al., 2007; P J Kennedy and Shapiro, 2009; Skaggs and McNaughton, 1998; Smith and Mizumori, 2006). Indeed, Keleman and Fenton (2010) showed that even in a single environment, the hippocampus can maintain two distinct maps and rapidly shift between them as needed to meet dynamic behavioral demands. Our findings are consistent with the idea that the hippocampus encodes contextual information, broadly defined to include spatial and non-spatial features of the situation (Smith and Bulkin, 2014). Indeed, part of the surprising stability of the trial and ITI representations could be related to the distinct and stable task requirements associated with each epoch.

The large scale multi-unit discharge at the trial start and trial end coincided with the transition between two distinct hippocampal representations, which raises the possibility that this burst of firing might serve to drive firing patterns out of one attractor space and into another (Rolls, 2007; Tom J Wills et al., 2005), pushing activity past a critical point to allow a shift in representational state (Steyn-Ross et al., 2010; Tkačik et al., 2015). Previous studies have identified individual neuronal responses near the start or end of trials (Ainge et al., 2007; Grieves et al., 2016; Hollup et al., 2001; Smith and Mizumori, 2006), but the impact of the response on population dynamics was only apparent when we examined the activity of large numbers of neurons. An intriguing possibility is that these bursts of firing may be involved in demarcating event boundaries. The hippocampus may play a critical role in event segmentation, the process of breaking continuous experience into discrete episodes (Zacks and Swallow, 2007). Hippocampal involvement in encoding the sequence of events (e.g. (Allen et al., 2016; Devito and Eichenbaum, 2011; Terada et al., 2017), for review see: (Davachi and DuBrow, 2015; Eichenbaum, 2014)) is consistent with an event segmentation explanation of ensemble activity, and recent work in humans found that an increase in activity in hippocampus is associated with event boundaries defined by change in distributed cortical representations (Baldassano et al., 2017). Event boundaries are often defined by a change in context, such as when a subject moves from one room to another or switches from one behavioral task to another (Horner et al., 2016), as was the case at the trial boundaries in the present study. We also found that a stronger representation of the trial start was associated with better performance, which is superficially similar to the finding that event segmentation in humans is linked to memory performance (Sargent et al., 2013; Zacks et al., 2006). However, this work has largely focused on the spontaneous segmentation of unique events, unlike the highly predictable and structured nature of our training trials so additional study will be needed to determine whether the multi-unit bursts we observed at trial boundaries are involved in more spontaneous forms of event segmentation.

## Materials and Methods

### Surgical and recording methods

Rats were surgically implanted with custom built moveable electrode arrays containing 16 insulated platinum iridium tetrodes, each composed of four 17 μm wires (California Fine Wire, Grover Beach, CA). Arrays were implanted with electrode tips located bilaterally just above the dorsal hippocampus (3.5mm posterior and 2.5mm lateral to bregma). Following recovery from surgery, the tetrodes were slowly lowered into the CA1 cell layer and rats began training on the behavioral task. Tetrodes were advanced over initial training and then left in place once rats reached asymptotic performance on the behavioral task. Multiunit recordings were sorted into constituent units using standard clustering techniques. We report on the activity of 2056 units recorded from 10 rats over 84 sessions (see Supplementary Table 1), counting individual recordings of units recorded over sessions (i.e. some units refer to a single neuron that was recorded on multiple sessions). Field potentials were sampled at 32kHz from one wire in each tetrode, and filtered between 0.1 and 6kHz, a representative signal was chosen from a tetrode located in the cell layer. All procedures complied with the guidelines established by the Cornell University Animal Care and Use Committee.

### Behavioral Procedure and Apparatus

Ten adult male Long-Evans rats were trained on a task designed to induce proactive mnemonic interference. Details on the task and the relationship between hippocampal activity and interference have been described elsewhere (Bulkin et al., 2016; Butterly et al., 2012; Law and Smith, 2012; Peters et al., 2013). Recordings took place in wooden chambers with a 60cm by 45cm floor and a removable divider (Supplementary Figure 1A). One side of the chamber served as an intertrial waiting area, the other contained two cups filled with odorized digging substrate. One of the cups was baited with a buried sugar pellet, reliably marked by odor, and rats learned to discriminate between 8 pairs of odors to retrieve rewards. On each trial, the divider was lifted, rats approached the cups and sampled odors, and dug for the sugar pellet. Rats were free to approach the baited cup first and completely ignore the unbaited odor (Supplementary Fig. 1B). Trials in which the rat sampled the unbaited odor and did not dig were marked as a correct rejection (Supplementary Fig. 1C) Trials were marked as errors if the rat dug in the unbaited cup (Supplementary Fig. 1D), any displacement of bedding was considered a digging response. Once the rat reached a behavioral criterion of 90% correct choices, a new set of odor pairs was presented, and training on this new set continued for 5 days. A subset of rats learned this new set in a distinct context. In the present paper, we focused on the responses within each session at the start and end of trials, the results of manipulating the context on behavioral performance and hippocampal ensemble activity have been described previously (Bulkin et al., 2016). Recordings were only taken on sessions with at least two units, although in most cases many more units were isolated: at least 10 units in 74/84 sessions (mean units/session 24.5; Supplementary Table 1 summarizes the number of units by rat/session).

### Data Analysis

#### Instantaneous Firing Rate

For each neuron, spike counts were binned across the entire session (100ms bins) and smoothed with a 5 bin moving average to construct a vector of instantaneous firing rate (IFR). This trace was normalized by subtracting the average and dividing by the standard deviation (i.e. z-scored) to produce a normalized IFR (IFRz). Because units showed a similar range of activity, analyses showed qualitatively similar results when using IFR or IFRz, but the latter prevented neurons with higher overall firing rates from dominating the analyses. Units with average rates greater than 4 spikes/second over the entire session were labeled as putative interneurons, and were not included in any of the analyses (235/2291 units were eliminated). Population vectors were defined as n x 1 vectors of IFRz at a given time, where n is the number of simultaneously recorded neurons.

The start and end of trials were identified as the moment the rat crossed an imaginary line corresponding to the location of the removable divider. Generally, rats only crossed this line once in each direction on each trial, but on those trials in which the rat entered the trial region and then returned back to the inter-trial waiting area only the first entry was used to mark the start of the trial. Trial start and end firing rate traces were calculated by linear interpolation of the IFR vector at times spanning +/-3s on each event. Spatial heatmaps were calculated by identifying the average firing rate of each neuron in 1.5cm square bins spanning the floor of the apparatus.

#### Local Field Potentials

The local field potential data was downsampled to 2kHz. The theta-delta ratio was identified as by filtering the signal (theta: 5-12Hz; delta: 2-4Hz) with a non-causal FIR filter, calculating RMS power with a 500ms sliding window, and taking the quotient.

#### Running Speed

Running speed was computed by applying a 1 second boxcar average to position, to eliminate spurious changes due to detection of the rats position or movement of the head. Average firing rate was calculated for binned running speeds (15 bins) separately for times near the event boundaries (+/- 1 second), in the trial (1 second after the start to 1 second before the end) and in the ITI (1 second after the trial end to 1 second before the trial start). A linear regression was calculated for data collected in each epoch to describe multi-unit activity as a function of running speed. The slope and intercept of these regressions were compared for neurons showing a slope that was significantly different from 0 (F-test, p<.01).

#### Generalized Linear Model

The strategy for applying a generalized linear model (GLM) that orthogonalized the contributions of spatial and temporal covariates was adapted from (Lepage et al., 2012; Truccolo et al., 2004). This approach relies on a geometry in the Fisher information of the GLM likelihood estimator to disambiguate activity due to a combination of multiple covariates. Applying this method to two-dimensional spatial data required a parameterization of spatial firing functions as activity depends on an interaction between the x and y co-ordinates defining the rat’s position. As such, we first described the both the temporal and spatial firing rate by fitting Gaussian curves and surfaces to the average firing data:

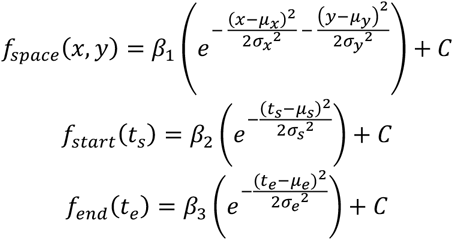

We modeled the coefficients *β_1_, β_2_, β_3_* by fitting a Poisson family GLM with a linear link function. Given the rat’s location (*x,y*), and the time relative to trial start (*t_s_*) and end (*t_e_*), the firing rate for each neuron was modeled as:

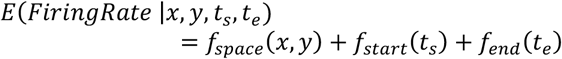

We also computed a model which included an additional factor for running speed:

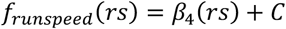

Because a linear link function in a Poisson GLM includes the possibility for negative rates, the *β_1_, β_2_, β_3,_ β_4_* parameters were restricted to be greater than or equal to 0. Since the true values could never be exactly zero, this does not break down the asymptotic orthogonalization results from (Lepage et al., 2012). The restriction of the parameter space means that parameters estimated to be 0 would have a negatively biased standard error in comparison to the traditional Wald-test and Likelihood Ratio Tests for generalized linear models. For this reason, we instead used a normal approximation significance test adapted from the normal theory bootstrap intervals given in (Efron and Tibshirani, 1994) to test if each parameter was significantly different than 0. If the bootstrapped sample for a parameter contained more than 10% of values selected exactly at 0, then a quantile-based significance test was used (Efron and Tibshirani, 1994). The quantile-based significance test was used in this case as the normality assumption on the bootstrapped sample no longer holds, however the quantile-based tests were not used across all observations to allow p-values to be calculated to more than 3 significant digits for significant parameters. The bootstrap was completed by randomly sampling 1 second blocks of data for each neuron with replacement. The blocking was done to account for the temporal dependencies in the data set (Gonçalves and Politis, 2011). Within each neuron, 250 bootstrap samples were created for each parameter in order to obtain a p-value. A significant effect was tabulated as any coefficient value with p<.01, for comparisons that grouped trial start and end responses together these p values were Bonferonni corrected.

#### Cross-Trial Distance Analysis

To compute cross-trial instantaneous ensemble firing rate similarity (Fig 3A-C), population vectors of instantaneous firing rate were assembled for times spanning +/-3seconds in 100ms intervals around each trial’s start and end. For each time point in each trial, the pairwise correlation between the associated population vector and all population vectors from all other trials at each time point was calculated. The average of these values was taken for each rat to form a cross trial correlation map for the session, each pixel representing the average correlation between population vectors taken from two time points across all pairs of trials. Correlation was calculated using both Pearson’s r and Kendall’s τ.

#### Classification of Trial versus ITI Responses

To identify the ability of neuronal populations to identify the current epoch, we trained linear discriminant classifiers to mark epoch based on activity. We first assembled population vectors spanning +/-3 seconds around the trial start and trial end, labeling each vector with the epoch in which it occurred. We took a random subset of half of these vectors and trained a linear discriminant classifier, separately for data occurring around the trial start or trial end, and tested the classifier on the remaining 50% of the data. This process was repeated 1000 times, on each iteration a different random subset was used to train/test the performance of the classifier. The overall performance of the classifier was measured as the median performance across iterations, and the 5th percentile of iterations was used to identify whether the classifier performed above chance.

#### Principal Component Analysis

To investigate the population dynamics of the event boundaries across the entire dataset, we created a synthetic dataset by sampling activity from defined time points around the trial start and end. This allowed visualization of activity in co-ordinates scaled by key sources of variance across a large population of neurons. One strategy for forming this synthetic dataset would be to randomly select activity from each neuron at a given time with respect to the event boundaries, however this approach would randomize covariance between neurons. Instead, we selected vectors from each session, preserving information about inter-neuron covariance when possible (i.e. within session) and randomizing when covariance data was unavailable (i.e. across sessions).

For each recording session (*ses*), we constructed population vectors (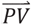) as the instantaneous firing rate at time points (*t*) spanning +/- 5s around each trial’s (*trial*) start and end in 100ms intervals.

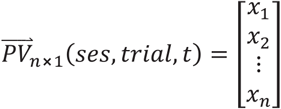

In the above equation, *x*_1_ indicates the firing rate of neuron 1 from recording session *ses,* trial number *trial* at the time specified by *t*. For instance, 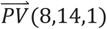 would contain the firing rates of all neurons recorded during session 8, on trial 14, 5.0 seconds before the trial start.

We then randomly selected a trial from each session and combined the population vectors across sessions (*v* = 84 total sessions), holding constant, to form pseudo-population vectors 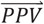.

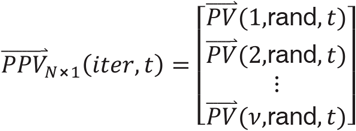

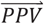 contains the firing rate of all *N* neurons (*N*=2056), on randomly selected trial, at some specific time (*t*) with respect to the trial start or end. It indicates what an ensemble of *N* neurons might look like at a given time.

We repeated this process over 200 iterations, and over all time windows, to form a ***N×M*** matrix. Each column of the matrix contains an iteratively selected ensemble firing rate at some time with respect to trial start or end.

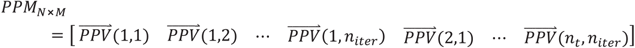

The total number of columns of *PPM*, denoted as *M*, is the product of the number of iterations (*n_iter_*) and the number of sampled time points (*n_t_*):

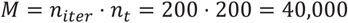

PC scores for the first 3 components of *PPM* were plotted directly, and a trajectory through PC space was calculated by applying the coefficients to produce these components back to the raw peri-event firing data.

An identical approach was taken for space, but here the grouping variable we used to combine vectors across sessions was the location of the rat associated with the instantaneous activity rather than the time of occurrence:

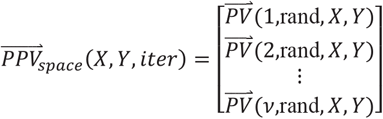

Rather than selecting randomly across trials, the spatial population vectors are selected from the set of firing rates associated with a specific (*X,Y*) location in space. *X* and *Y* were bins that spanned the range of the recording apparatus, in 20 pixel (about 3cm) square bins. After eliminating bins that were not visited by all of the rats in the experiment, 208 spatial bins remained, producing a spatial pseudo-population matrix (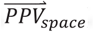) with a similar size as the one used in the temporal analysis:

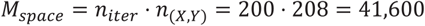

## Acknowledgments

We thank Leela Patel, Carly Britton, and Luke Grosvenor for assistance with animal training and electrode hyperdrive fabrication. We also would like to thank Khena Swallow, Sam Whitehead and Vikram Gadagkar for feedback on the analyses and development of ideas presented in the manuscript. This work was supported by an NIH grant R01 MH083809 to D.M.S.

## References

Ainge, J.A., Tamosiunaite, M., Woergoetter, F., Dudchenko, P.A., 2007. Hippocampal CA1 Place Cells Encode Intended Destination on a Maze with Multiple Choice Points. J. Neurosci. 27, 9769–9779. doi:10.1523/JNEUROSCI.2011-07.2007

Allen, T.A., Salz, D.M., McKenzie, S., Fortin, N.J., 2016. Nonspatial Sequence Coding in CA1 Neurons. J. Neurosci. 36, 1547–1563. doi:10.1523/JNEUROSCI.2874-15.2016

Alme, C.B., Miao, C., Jezek, K., Treves, A., Moser, E.I., Moser, M.-B., 2014. Place cells in the hippocampus: eleven maps for eleven rooms. Proc. Natl. Acad. Sci. U. S. A. 111, 18428–35. doi:10.1073/pnas.1421056111

Anderson, M.I., Jeffery, K.J., 2003. Heterogeneous modulation of place cell firing by changes in context. J. Neurosci. 23, 8827–35.

Baldassano, C., Chen, J., Zadbood, A., Pillow, J.W., Hasson, U., Norman, K.A., 2017. Discovering Event Structure in Continuous Narrative Perception and Memory. Neuron 95, 709–721.e5. doi:10.1016/j.neuron.2017.06.041

Bulkin, D.A., Law, L.M., Smith, D.M., 2016. Placing memories in context: Hippocampal representations promote retrieval of appropriate memories. Hippocampus 26, 958–971. doi:10.1002/hipo.22579

Butterly, D.A., Petroccione, M.A., Smith, D.M., 2012. Hippocampal context processing is critical for interference free recall of odor memories in rats. Hippocampus 22, 906–913. doi:10.1002/hipo.20953

Buzsáki, G., Horváth, Z., Urioste, R., Hetke, J., Wise, K., 1992. High-frequency network oscillation in the hippocampus. Science 256, 1025–7.

Colgin, L.L., Moser, E.I., Moser, M.B., 2008. Understanding memory through hippocampal remapping. Trends Neurosci 31, 469–477. doi:10.1016/j.tins.2008.06.008

Davachi, L., DuBrow, S., 2015. How the hippocampus preserves order: the role of prediction and context. Trends Cogn. Sci. 19, 92–99. doi:10.1016/j.tics.2014.12.004

Devito, L.M., Eichenbaum, H., 2011. Memory for the order of events in specific sequences: contributions of the hippocampus and medial prefrontal cortex. J. Neurosci. 31, 3169–75. doi:10.1523/JNEUROSCI.4202-10.2011

Efron, B., Tibshirani, R., 1994. An introduction to the bootstrap. Chapman & Hall.

Eichenbaum, H., 2014. Time cells in the hippocampus: a new dimension for mapping memories. Nat. Rev. Neurosci. 15, 732–744. doi:10.1038/nrn3827

Eichenbaum, H., Kuperstein, M., Fagan, A., Nagode, J., 1987. Cue-sampling and goal-approach correlates of hippocampal unit activity in rats performing an odor-discrimination task. J Neurosci 7, 716–732.

Eschenko, O., Mizumori, S.J.Y., 2007. Memory influences on hippocampal and striatal neural codes: Effects of a shift between task rules. Neurobiol. Learn. Mem. 87, 495–509. doi:10.1016/j.nlm.2006.09.008

Ferbinteanu, J., Shapiro, M.L., 2003. Prospective and retrospective memory coding in the hippocampus. Neuron 40, 1227–39.

Gonçalves, S., Politis, D., 2011. Discussion: Bootstrap methods for dependent data: A review. J. Korean Stat. Soc. doi:10.1016/j.jkss.2011.07.003

Grieves, R.M., Wood, E.R., Dudchenko, P.A., 2016. Place cells on a maze encode routes rather than destinations. Elife 5. doi:10.7554/eLife.15986

Griffin, A.L., Eichenbaum, H., Hasselmo, M.E., 2007. Spatial Representations of Hippocampal CA1 Neurons Are Modulated by Behavioral Context in a Hippocampus-Dependent Memory Task. J. Neurosci. 27, 2416–2423. doi:10.1523/JNEUROSCI.4083-06.2007

Holland, P.C., Bouton, M.E., 1999. Hippocampus and context in classical conditioning. Curr. Opin. Neurobiol. 9, 195–202. doi:10.1016/S0959- 4388(99)80027-0

Hollup, S.A., Molden, S., Donnett, J.G., Moser, M.B., Moser, E.I., 2001. Accumulation of hippocampal place fields at the goal location in an annular watermaze task. J. Neurosci. 21, 1635–44.

Horner, A.J., Bisby, J.A., Wang, A., Bogus, K., Burgess, N., 2016. The role of spatial boundaries in shaping long-term event representations. Cognition 154, 151–164. doi:10.1016/j.cognition.2016.05.013

Jezek, K., Henriksen, E.J., Treves, A., Moser, E.I., Moser, M.B., 2011. Theta-paced flickering between place-cell maps in the hippocampus. Nature 478, 246–249. doi:10.1038/nature10439

Kelemen, E., Fenton, A.A., 2010. Dynamic grouping of hippocampal neural activity during cognitive control of two spatial frames. PLoS Biol 8, e1000403. doi:10.1371/journal.pbio.1000403

Kennedy, P.J., Shapiro, M.L., 2009. Motivational states activate distinct hippocampal representations to guide goal-directed behaviors. Proc. Natl. Acad. Sci. 106, 10805–10810. doi:10.1073/pnas.0903259106

Kennedy, P.J., Shapiro, M.L., 2009. Motivational states activate distinct hippocampal representations to guide goal-directed behaviors. Proc Natl Acad Sci U S A 106, 10805–10810. doi:10.1073/pnas.0903259106

Law, L.M., Smith, D.M., 2012. The anterior thalamus is critical for overcoming interference in a context-dependent odor discrimination task. Behav Neurosci 126, 710–719. doi:10.1037/a0029698

Lepage, K.Q., MacDonald, C.J., Eichenbaum, H., Eden, U.T., 2012. The statistical analysis of partially confounded covariates important to neural spiking. J. Neurosci. Methods 205, 295–304. doi:10.1016/j.jneumeth.2011.12.021

Leutgeb, J.K., Leutgeb, S., Moser, M.-B., Moser, E.I., 2007. Pattern separation in the dentate gyrus and CA3 of the hippocampus. Science 315, 961–6. doi:10.1126/science.1135801

Leutgeb, S., Leutgeb, J.K., Barnes, C.A., Moser, E.I., McNaughton, B.L., Moser, M.B., 2005. Independent codes for spatial and episodic memory in hippocampal neuronal ensembles. Science (80-.). 309, 619–623. doi:10.1126/science.1114037

MacDonald, C.J., Lepage, K.Q., Eden, U.T., Eichenbaum, H., 2011. Hippocampal “time cells” bridge the gap in memory for discontiguous events. Neuron 71, 737–49. doi:10.1016/j.neuron.2011.07.012

Muller, R.U., Kubie, J.L., 1987. The effects of changes in the environment on the spatial firing of hippocampal complex-spike cells. J. Neurosci. 7, 1951–68.

Muzzio, I.A., Levita, L., Kulkarni, J., Monaco, J., Kentros, C., Stead, M., Abbott, L.F., Kandel, E.R., 2009. Attention enhances the retrieval and stability of visuospatial and olfactory representations in the dorsal hippocampus. PLoS Biol 7, e1000140. doi:10.1371/journal.pbio.1000140

Nadel, L., 2008. The Hippocampus and Context Revisited, in: Mizumori, S.J. (Ed.), Hippocampal Place Fields: Relevance to Learning and Memory. Oxford University Press, New York, pp. 3–15.

Neymotin, S.A., Talbot, Z.N., Jung, J.Q., Fenton, A.A., Lytton, W.W., 2017. Tracking recurrence of correlation structure in neuronal recordings. J. Neurosci. Methods 275, 1–9. doi:10.1016/j.jneumeth.2016.10.009

Peters, G.J., David, C.N., Marcus, M.D., Smith, D.M., 2013. The medial prefrontal cortex is critical for memory retrieval and resolving interference. Learn Mem 20, 201–209. doi:10.1101/lm.029249.112

Rolls, E.T., 2007. An attractor network in the hippocampus: theory and neurophysiology. Learn. Mem. 14, 714–31. doi:10.1101/lm.631207

Rudy, J.W., 2009. Context representations, context functions, and the parahippocampal-hippocampal system. Learn. Mem. 16, 573–85. doi:10.1101/lm.1494409

Sargent, J.Q., Zacks, J.M., Hambrick, D.Z., Zacks, R.T., Kurby, C.A., Bailey, H.R., Eisenberg, M.L., Beck, T.M., 2013. Event segmentation ability uniquely predicts event memory. Cognition 129, 241–255. doi:10.1016/j.cognition.2013.07.002

Schlesiger, M.I., Boublil, B.L., Hales, J.B., Leutgeb, J.K., Leutgeb, S., 2018. Hippocampal Global Remapping Can Occur without Input from the Medial Entorhinal Cortex. Cell Rep. 22, 3152–3159. doi:10.1016/j.celrep.2018.02.082

Skaggs, W.E., McNaughton, B.L., 1998. Spatial firing properties of hippocampal CA1 populations in an environment containing two visually identical regions. J. Neurosci. 18, 8455–66.

Smith, D.M., Bulkin, D.A., 2014. The form and function of hippocampal context representations. Neurosci Biobehav Rev 40C, 52–61. doi:10.1016/j.neubiorev.2014.01.005

Smith, D.M., Mizumori, S.J., 2006. Learning-related development of context-specific neuronal responses to places and events: the hippocampal role in context processing. J Neurosci 26, 3154–3163. doi:10.1523/JNEUROSCI.3234-05.2006

Steyn-Ross, D.A., Steyn-Ross, M.L., Wilson, M.T., Sleigh, J.W., 2010. Phase transitions in single neurons and neural populations: Critical slowing, anesthesia, and sleep cycles, in: Modeling Phase Transitions in the Brain. Springer New York, New York, NY, pp. 1–26. doi:10.1007/978-1-4419-0796-7_1

Terada, S., Sakurai, Y., Nakahara, H., Fujisawa, S., 2017. Temporal and Rate Coding for Discrete Event Sequences in the Hippocampus. Neuron 94, 1248–1262.e4. doi:10.1016/j.neuron.2017.05.024

Terrazas, A., Krause, M., Lipa, P., Gothard, K.M., Barnes, C.A., McNaughton, B.L., 2005. Self-Motion and the Hippocampal Spatial Metric. J. Neurosci. 25, 8085–8096. doi:10.1523/JNEUROSCI.0693-05.2005

Tkačik, G., Mora, T., Marre, O., Amodei, D., Palmer, S.E., Berry, M.J., Bialek, W., 2015. Thermodynamics and signatures of criticality in a network of neurons. Proc. Natl. Acad. Sci. U. S. A. 112, 11508–13. doi:10.1073/pnas.1514188112

Truccolo, W., Eden, U.T., Fellows, M.R., Donoghue, J.P., Brown, E.N., 2004. A Point Process Framework for Relating Neural Spiking Activity to Spiking History, Neural Ensemble, and Extrinsic Covariate Effects. J. Neurophysiol. 93, 1074–1089. doi:10.1152/jn.00697.2004

Wills, T.J., Lever, C., Cacucci, F., Burgess, N., O’Keefe, J., 2005. Attractor Dynamics in the Hippocampal Representation of the Local Environment. Science (80-.). 308, 873–876. doi:10.1126/science.1108905

Wills, T.J., Lever, C., Cacucci, F., Burgess, N., O’Keefe, J., 2005. Attractor dynamics in the hippocampal representation of the local environment. Science 308, 873–6. doi:10.1126/science.1108905

Wood, E.R., Dudchenko, P.A., Robitsek, R.J., Eichenbaum, H., 2000. Hippocampal neurons encode information about different types of memory episodes occurring in the same location. Neuron 27, 623–633.

Zacks, J.M., Speer, N.K., Vettel, J.M., Jacoby, L.L., 2006. Event understanding and memory in healthy aging and dementia of the Alzheimer type. Psychol. Aging 21, 466–482. doi:10.1037/0882-7974.21.3.466

Zacks, J.M., Swallow, K.M., 2007. Event Segmentation. Curr. Dir. Psychol. Sci. 16, 80–84. doi:10.1111/j.1467-8721.2007.00480.x

